# Life on leaves : uncovering temporal dynamics in Arabidopsis’ leaf microbiota

**DOI:** 10.1101/2021.07.06.450897

**Authors:** Juliana Almario, Maryam Mahmudi, Samuel Kroll, Mathew Agler, Aleksandra Placzek, Alfredo Mari, Eric Kemen

## Abstract

Leaves are primarily responsible for the plant’s photosynthetic activity. Thus, changes in the phyllosphere microbiota, which includes deleterious and beneficial microbes, can have far reaching effects on plant fitness and productivity. In this context, identifying the processes and microorganisms that drive the changes in the leaf microbiota over a plant’s lifetime is crucial. In this study we analyzed the temporal dynamics in the leaf microbiota of *Arabidopsis thaliana*, integrating both compositional changes and changes in microbe-microbe interactions via the study of microbial networks. Field-grown *Arabidopsis* were used to follow leaf bacterial, fungal and oomycete communities, throughout the plant’s growing season (extending from November to March), over three consecutive years. Our results revealed the existence of conserved time patterns, with microbial communities and networks going through a stabilization phase (decreasing diversity and variability) at the beginning of the plant’s growing season. Despite a high turnover in these communities, we identified 19 ‘core’ taxa persisting in Arabidopsis leaves across time and plant generations. With the hypothesis these microbes could be playing key roles in the structuring of leaf microbial communities, we conducted a time-informed microbial network analysis which showed core taxa are not necessarily highly connected network ‘hubs’ and ‘hubs’ alternate with time. Our study shows that leaf microbial communities exhibit reproducible dynamics and patterns, suggesting it could be possible to predict and drive these microbial communities to desired states.

## Introduction

Leaves are primarily responsible for the plant’s photosynthetic activity and gaseous exchange. Consequently, leaf health and performance have a direct effect on plant growth and fitness [1]. Leaves are colonized by a wide range of microbes, including bacteria, archaea, and microeukaryotes like fungi and oomycetes. While natural openings on leaves such as stomata, hydathodes or wounds represent entry points for major plant-pathogens, they also house commensal and even beneficial microbes, leading to plant-protecting effects [2, 3]. There is increasing interest particularly by plant breeders in microbiota-engineering approaches to promote the growth and health of crops through beneficial microbes [4]. In this context, the understanding of the processes that shape the composition of leaf microbiota is an essential step.

There is a level of specificity between plants and their leaf microbiota as studies have repeatedly shown that different plant lineages tend to harbor quantitatively different microbial consortia in their leaves [5], with differences even observed between ecotypes of the same plant species [6]. Although it is unclear how plants can selectively recruit certain microbial groups, the soil in which plants grow appears to be an important driver [6, 7]. The study of plant microbiota over different developmental stages suggests that as the plant grows, the microbiota becomes more tissue-specific with major differences observed between root and shoot microbiota [8]. There is increasing awareness of the fact that plant-associated microbiota are not static but dynamic communities changing through time and shaped convergently by environmental and host cues. Recent studies have followed the dynamics of microbiota formation in leaves [9–12] and roots [13] but few of them have conducted a cross kingdom survey, integrating both bacterial and micro-eukaryotic communities.

Microbe-microbe interactions such as mutualism, antagonism or predation, shape the composition of microbial communities. Correlation network analyses on the abundance of microbial taxa, can be used to infer microbial interactions in a community. The study of microbial networks over time can inform us about the dynamics of these interactions and how they relate to changes in the diversity and structure of microbial communities [14]. Yet, such approaches have rarely been applied to investigate how plant-associated microbiota change through the plant’s life.

Given the complexity of leaf microbial communities, assigning ecological roles and ecological importance to individual taxa is extremely challenging. Concepts based on the persistence of a microbe (core taxa) and/or its importance on microbial networks (hubs taxa) have been applied to identify microorganisms playing key roles in leaf communities [15, 16]. Although the large majority of leaf microbes show scattered distributions with highly-fluctuating occurrences in plant leaves across environments and time, some microorganisms achieve a stable presence in plant populations [17]. It is unclear how these “core” microbes are able to systematically colonize the host-plant, but it could involve re-colonization processes [18] or vertical inheritance *via* seeds [19]. The stability of the associations between “core” microbes and the host-plant suggests a high level of adaptation to the leaf niche on the microbe side. This can involve traits associated to plant-colonization and infection as suggested for leaf pathogenic *Pseudomonas viridiflava* [17], but it can also involve the capacity of the microorganism to re-shape the leaf microbiota, as part of a ‘niche construction’ strategy. Notably, Agler *et al*. (2016) [16] showed that the inoculation of the leaf-pathogenic oomycete Albugo on Arabidopsis plants translates into decreased microbial diversity on leaves and altered microbiota profiles. The analysis of microbial interaction networks in the leaf microbiota showed Albugo acts as a network ‘hub’, showing the highest level of connections (interactions) with other microbes, which would allow it to influence the structure of the leaf microbiota. Because of its hub characteristics and experimentally proven impact on leaf microbial communities it has been proposed as a ‘keystone’ taxon of the leaf microbiota in Arabidopsis. However, it is still unclear whether re-shaping the leaf microbiome contributes to persistence of core taxa.

The aim of this study was to analyze the temporal dynamics in the leaf microbiota of Arabidopsis, integrating both compositional changes and changes in microbe-microbe interactions via the study of microbial networks. Amplicon sequencing was used to follow leaf bacterial, fungal and oomycete communities in a field experiment throughout the plant’s growing season (extending from November to March). The experiment was carried out over three consecutive years in order to capture long-term dynamics. Our results reveal seasonal/monthly patterns associated with reproducible changes in particular groups across kingdoms like Sphingomonadales and Actinomycetales bacteria, Microbotryales and Sporidiobolales fungi and Peronosporales oomycetes. Despite a high level of stochasticity in microbial colonization of the leaf, we identified 19 taxa that were consistently present (core taxa), including putative pathogenic and beneficial taxa. Between November and February, the diversity and variability of leaf microbial communities decreased, as microbial networks stabilized (changed less) and exhibited decreasing complexity (number of nodes and connections). With the hypothesis that certain microbes play a predominant role in the structuring and stability of these communities, we focused on the identification of microbes having both a persistent presence in Arabidopsis leaves (core microbes) and a high connectivity in leaf microbiota networks (hub microbes).

## Material and Methods

### Common garden experiment

To study the temporal dynamics of *A. thaliana*’s leaf microbiota, we conducted a common garden experiment where *A. thaliana* plants were sampled every month from November to March, covering the plant’s natural growing season, including the vegetative and early reproductive growth phases (Fig. 1). The experiment was conducted as described in Agler *et al*., 2016 [16]. Briefly, surface-sterilized seeds were germinated on Jiffy pellets for 10 days under greenhouse conditions, before transferring to the field. To take into account host genetic variability, four global *Arabidopsis thaliana* ecotypes were used (Ws-0, Col-0, Ksk-1 and Sf-2). The field was divided into nine experimental plots which were planted with 10 plants per ecotype, in a randomized set-up. At each sampling point, whole leaf samples were taken from 2-4 randomly selected plants per ecotype. The whole experiment was repeated three times in 2014-2015, 2015-2016 and 2016-2017. The field is located at the Max-Planck Institute for Plant Breeding Research (Cologne, Germany) (Supplementary Table 1).

**Figure 1.**
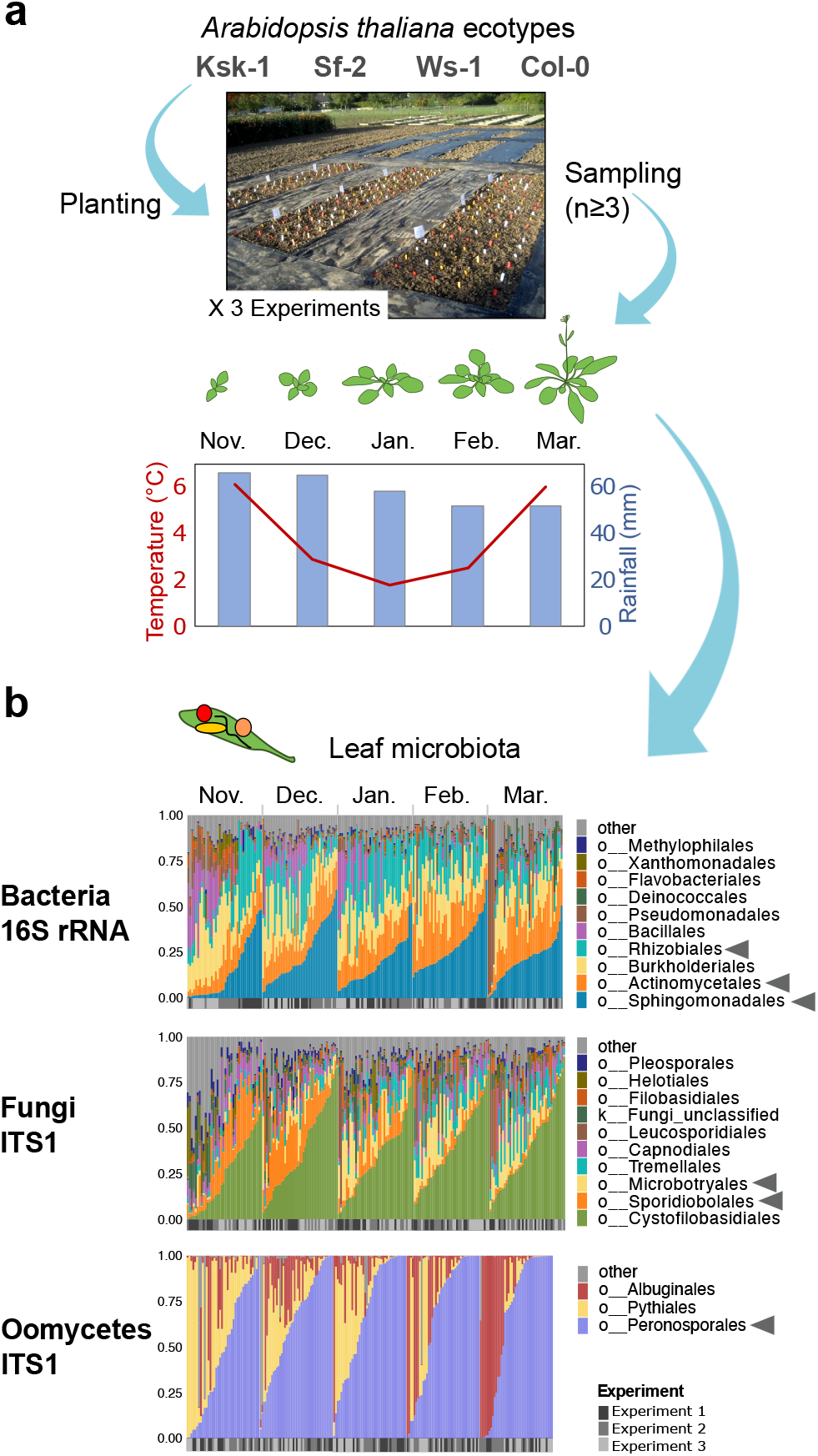
Following leaf microbiota changes throughout *A. thaliana’s* growing season. **(A)** Experimental set-up. Four global *Arabidopsis* accessions were planted in a common garden (Max Planck institute, Cologne, Germany). Every month from November to March, 3 plant individuals per ecotype were collected and leaf samples were taken for microbiota analysis. The experiment was repeated three times over years 2014-2015 (experiment 1), 2015-2016 (2) and 2016-2017 (3), with a total number of 206 plant leaf samples analyzed (see Supplementary Table 1). **(B)** Composition of the leaf microbiota. Microbiota analysis was conducted via Illumina-based amplicon sequencing (Miseq 2 x 300 bases). Taxonomic markers included the bacterial 16S rRNA v5-v7 region, fungal ITS1 and the oomycete ITS1 region. Histograms show the relative abundance of the main microbial groups (order level) in single samples aggregated by ‘month’. Grey boxes below histograms indicate the ‘experiment’. Arrowheads indicate taxa exhibiting marked seasonal patterns (see Fig. S2).

### DNA extraction and amplicon sequencing

Samples were processed exactly as described in Agler et al., 2016 [16]. Briefly, whole leaf samples were crushed and used for phenolchloroform-based DNA extraction. The obtained DNA was used for two-step PCR amplification of the V5-V7 region of the bacterial 16S rRNA (primers B799F/B1194R), the fungal ITS1 region (primers ITS1F/ITS2) and the oomycetes ITS1 region (primers ITS1O/5.8s-O-R). Blocking oligos were used to reduce plant DNA amplification [20]. Purified PCR products were pooled in equimolar amounts before sequencing on three Illumina MiSeq runs (2 x 300 bp reads) with 10% PhiX control. Primers targeting the oomycete ITS1 region also produced “non-oomycete” reads but at a very marginal level (3%).

### Amplicon sequencing data analysis

Amplicon sequencing data was processed in Mothur [21] as described in Karasov *et al*., 2018 [17]. Single-end reads were paired (*make.contigs* command) and paired reads with more than 5 bases overlap between the forward and reverse reads were kept. Only 100-600 bases long reads were retained (*screen.seqs*). Chimeras were checked using Uchime in Mothur with more abundant sequences as reference (*chimera.uchime*, abskew = 1.9). Sequences were clustered into OTUs at the 97% similarity threshold using the VSEARCH program in Mothur (*cluster*, dgc method). Individual sequences were taxonomically classified using the rdp classifier method (*classify.seqs*, consensus confidence threshold set to 80) and the greengenes database (13_8 release) for 16S rRNA data, the UNITE_public database (version 12_2017) for fungal ITS1 and the Pr2 (version 4.10.0) for oomycete ITS1. The PhiX genome was included in each of the databases to improve the detection of remaining PhiX reads. Each OTU was then taxonomically classified (*classify.otu*, consensus confidence threshold set to 66), OTUs with unknown taxonomy at the Kingdom level were removed, as were low abundance OTUs (< 50 reads, *split.abund*).

Sample alpha-diversity analysis was conducted on OTU abundance tables, using Shannon’s H diversity index (estimate_richness function in phyloseq package). Data normality was checked (Shapiro-Wilk’s test) and means were compared by ANOVA followed by Tukey’s HSD (*P* < 0.05). Beta-diversity analyses were conducted on transformed (log_10_ (*x* + 1)) OTU relative abundance tables. Bray-Curtis dissimilarities between samples were computed and used for non-metric multidimensional scaling ordination (NMDS, function ‘ordinate’, Phyloseq package). A PerMANOVA analysis on Bray-Curtis dissimilarities was conducted to identify the main factors influencing the structure of the leaf microbiota (‘Adonis’, Vegan package, 10 000 permutations, *P* < 0.05, explanatory categorical variables: Experiment x Month x Ecotype). A beta-dispersion analysis on Bray-Curtis dissimilarities was conducted to compare sample-to-sample variation within each month of sampling (multivariate homogeneity of group dispersions analysis, ‘betadisper’, Vegan package). Differences between conditions were tested using a non-parametric multivariate test (Dunn’s test, *P* < 0.05). All analyses were conducted in R 3.6.1.

### Identification of a core leaf microbiota in A. thaliana

Core taxa were identified as OTUs showing high-occurrence over time (≥ 95% for fungi and oomycete, ≥ 98% for bacteria) in each of the three experiments analyzed. A higher cut-off was used for bacteria (98% occurrence) as they exhibited a higher average occurrence compared to fungi and oomycetes. The taxonomical classification of core OTUs was used to compute pairwise dissimilarities (distances) between OTUs (‘daisy’ function, Cluster package in R, Gower’s distance) which were used for hierarchical clustering (‘hclust’ function, Cluster package in R). The obtained dendrogram was modified in the browser version of iTOL (version 5.5.1) [22].

### Network analysis

Bacteria, fungi and oomycete OTU tables were merged and used for correlations calculation using either the Spearman correlation coefficient in Co-Net [23] or the SparCC algorithm [24] which relies on Aitchison’s log-ratio analysis and is designed to deal with compositional data with high sparsity like this data set (sparsity = 74%) [25]. OTU tables were filtered to OTUs present in at least 5 samples with >10 reads per OTU (sparsity = 53%). For the Co-Net based analysis, OTUs relative abundances were calculated and the obtained OTU tables were transformed (log_10_ (*x* +1)) before calculating Spearman correlation scores using Co-Net in Cytospace [26]. Parameters included the selection of top 5% correlations (edge selection, quantile=0.05, top and bottom) and the computing of *P*-values by Fisher’s Z-score with multiple-test correction (Bonferroni, *P*-value = 0.001). For the SparCC based analysis, the filtered OTU tables (OTU raw abundances) were used to calculate SparCC correlation scores (with default parameters). Pseudo *P*-values were inferred from 1000 bootstraps. Only correlations with *P* < 0.001 were kept for further analyses. Cytoscape (version 3.7.1) was used for network visualization and determination of betweenness centrality (i.e. the fraction of shortest paths passing through a given node) and closeness centrality values (i.e. the average shortest distance from given node to each other node). Node-rewiring score (Dn-score) was calculated via the DyNet package in Cytoscape [27]. For each node, its connected neighbors are compared between two networks and the changes (rewiring) are quantified. Microbial hubs were identified as top 5% OTUs showing maximum betweenness centrality and closeness centrality scores.

Sequencing data is available under NCBI Bioproject PRJNA438596. OTU tables and scripts are available here https://github.com/IshtarMM/Dynamic_LeafMicrobiome

## Results

### The leaf microbiota is highly dynamic

To study temporal dynamics in the leaf microbiota, we grew four *A. thaliana* ecotypes in a common garden and surveyed the changes in their leaf microbiota via amplicon-sequencing (bacteria, fungi and oomycetes). Leaf samples were taken monthly between November and March (5 months), thus covering most of the plant’s growing season over autumn and winter (Fig. 1). To identify the main factors shaping leaf microbial communities we conducted multivariate analyses including non-metric multidimensional scaling (NMDS; Fig. S1A) and Permutational multivariate ANOVA (Bray-Curtis dissimilarities, *P* < 0.05; Fig. S1B) on the relative abundance of bacterial, fungal and oomyceyte taxa (OTUs defined at 97% similarity). These analyses showed a marginal effect of the plant ecotype (2-4% explained variance) but an important effect of the time of sampling (32-40% explained variance; factors ‘Month’, ‘Experiment’, and their interaction; Fig. S1B), confirming leaf microbial communities are highly variable in time (i.e. dynamic). Although variability between experiments was significant (4-13% explained variance), the ‘Month’ of sampling was an important factor (11-15% explained variance; Fig. S1 B, Fig. S1A), suggesting the existence of seasonal/monthly patterns in these microbial communities. Such patterns were easily observable when considering changes in the relative abundance of high-abundance microbial orders (Fig. 1). For example, the abundance of Sphingomonadales and Actinomyceteales increased throughout the plant’s growing season, while the abundance of Rhizobiales tended to decrease. As for fungi, the abundance of Microbotryales increased while that of Sporidiobolales decreased. Interestingly, the abundance of Peronosporales oomycetes, which include *A. thaliana*’s pathogen *Hyaloperonospora* spp., increased with time, reaching maximum values at the end of the plant’s growing season (Tukey HSD test, *P* < 0.05) (Fig. 1; Fig. S2).

### Persistent (core) taxa in the leaf microbiota

We aimed to identify microbial groups showing a persistent presence throughout the plant’s life, hypothesizing they might play important roles in plant-microbe and microbe-microbe interactions within the microbiota. Highly persistent microbes (≥ 95% sample occurrence for fungi and oomycete, ≥ 98% for bacteria) varied considerably between experiments with only 19 out of 67 OTUs (28%) showing robust patterns across experiments (Fig. S3). Notably, these persistent core taxa (1 oomycete, 6 fungi and 12 bacteria OTUs) included known Arabidopsis pathogens like the obligate biotrophic oomycete *Hyaloperonospora* sp. (Otu00001) as well as bacterial taxa known to colonize Arabidopsis leaves, including Sphingomonas sp. (OTUs), Methylobacterium sp (Otu000002), and Variovorax (Otu000010). Persistent fungal taxa included two ascomycetes (Cladosporium sp. Otu00004 and Otu00012) and four basidiomycete yeast (Dioszegia sp. Otu00013, Itersonilia sp. Otu00005, Sporidiobolus sp. Otu00002, and Udeniomyces sp. Otu00001) (Fig. 2A). The relative abundance of these core taxa changed throughout the plant’s growing season reaching a maximum in February where it represented as much as 49, 52 and 71% of the bacterial, fungal and oomycete communities, respectively (Fig. 2B). These results indicate that despite the highly dynamic and stochastic nature of the leaf microbiota, a limited number of microbes -- only 19 out of 3058 OTUs (0.62%) -- consistently co-colonize plant leaves. This suggests a high degree of adaptation to this niche but also frequent interactions with one another.

**Figure 2.**
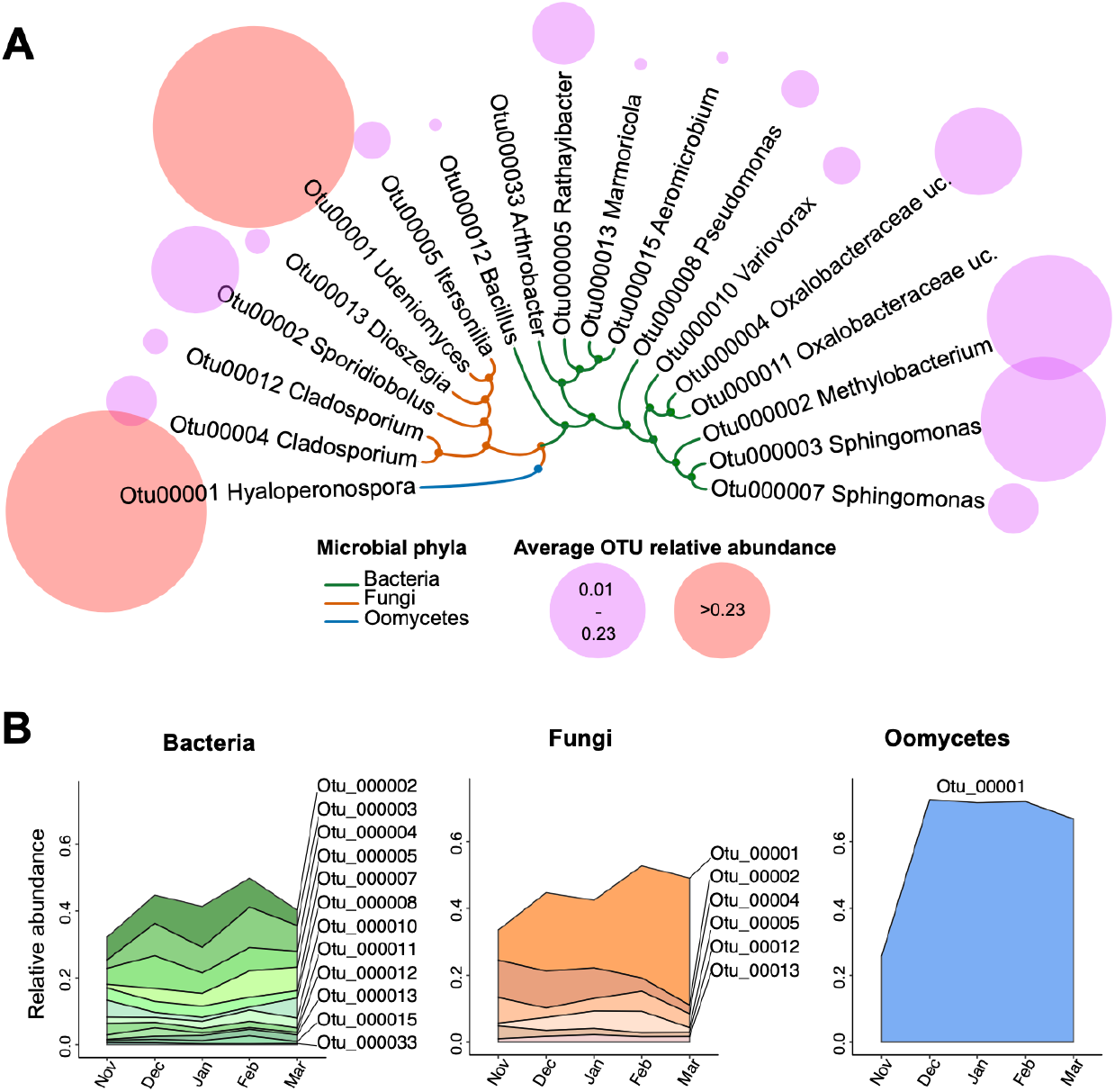
Persistent core members of the Arabidopsis leaf microbiota. **(A)** Core taxa were identified as OTUs showing high-occurrence (≥ 95% for fungi and oomycete, ≥ 98% for bacteria) in each of the three experiments. Bubbles depict the average relative abundance of each core OTU, per sample. The dendrogram depicts taxonomical distances between OTUs (hierarchical-clustering on Gower distances from OTU taxonomy). **(B)** Changes in the relative abundance of core taxa over time (month averages; n > 38 samples per month).

Diversity and variability of the leaf microbiota decrease throughout the plant’s growing season as communities stabilize. With the hypothesis that leaf-associated microbial communities become increasingly stable throughout the plant’s growing season, we analyzed their dynamics in terms of alpha-diversity (number of taxa in the community), within-month variability (plant-to-plant differences in community composition) and variability between consecutive months (month-to-month differences in community composition). While bacterial alpha-diversity (Shannon’s H index) remained unchanged, fungal and oomycete alpha-diversity decreased with significant differences observed between November and the last two months, February and March (ANOVA followed by Tukey’s HSD, P < 0.05) (Fig. 3). A similar trend was observed for within-month variability (sample distance to the group centroid), as variability of bacterial and fungal communities decreased progressively from November to February (Dunn test, P < 0.05) (Fig. 3). Similarly, a progressive decrease in between-month variability (sample-to-sample distances between consecutive months) was observed for bacterial and fungal communities (Dunn test, P < 0.05) (Fig. 3). Oomycete communities exhibited similar trends but the dynamics were less pronounced due to higher data variability. Together these results suggest that throughout the plant’s growing season, leaf microbial communities become progressively less diverse, more similar between plant individuals, and less variable in time. This suggests leaf communities go through a consolidation and stabilization phase from November to February.

**Figure 3.**
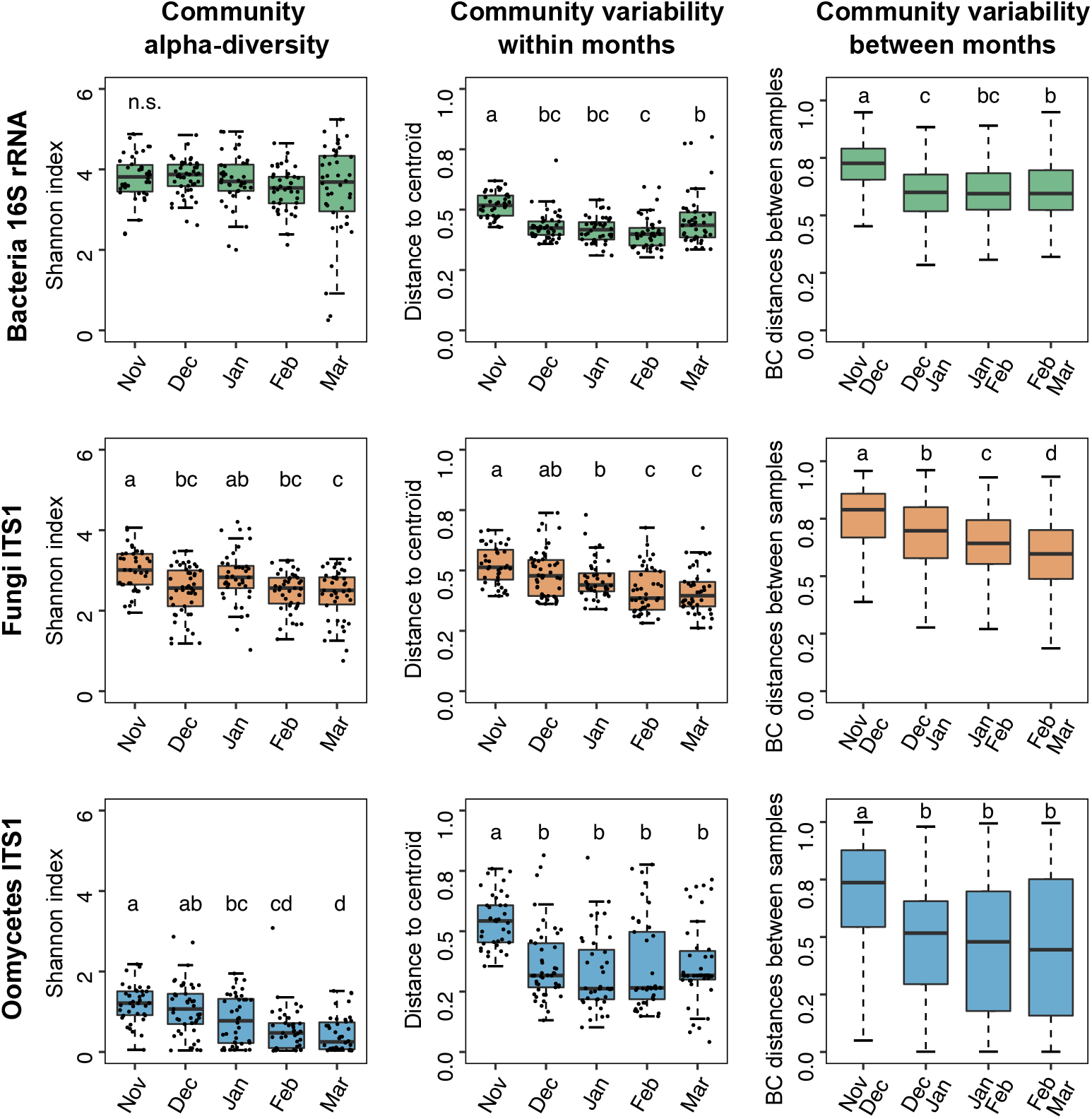
Changes in alpha-diversity and variability in leaf microbial communities, over time. Alpha diversity (Shannon’s H index), within-month variability (distance to the group centroid; beta-dispersion) and between-month variability (Bray-Curtis distances between samples from consecutive months) in bacterial, fungal and oomycete communities. Each box-plot shows combined data from the 3 experiments with n>38 samples per month. Dots represent individual samples, whiskers depict the dispersion of the data (1.5 x interquartile range) and different letters indicate significant differences between groups (Shannon index: ANOVA followed by Tukey’s HSD, *P* < 0.05; distances: Dunn test, *P* < 0.05). Single BC distances between samples are not shown because of the high number of comparisons (>700).

### Interaction networks within the leaf microbiota stabilize over time

Microbial networks computed from correlation of species abundances, are used to infer potential interactions between microbes within a community. To determine if/how leaf microbial networks changed over time, we used taxa abundance data from each time point (month) to generate five ‘month’ networks (Fig. 4A). Because the data was highly sparse (53% sparsity), the SparCC algorithm (optimized for sparse data) was used for network calculation [25]. The five networks differed in terms of general characteristics like the number of nodes (number of taxa) and edges (correlations between taxa; syn. connections) with no clear pattern, except for the month of ‘February’ which had both the lowest number of nodes and the lowest number of edges (Fig. 4B). Similarly, the nodes of this network had the lowest number of interactions (node degree), going from 70 on average in January to only 10 on average in February (Fig. 4BC; Dunn test, *P* < 0.05). This confirmed that microbial networks indeed change throughout the plant’s growing season and suggested major restructuring events around the month of February, when the network exhibited minimal complexity.

**Figure 4.**
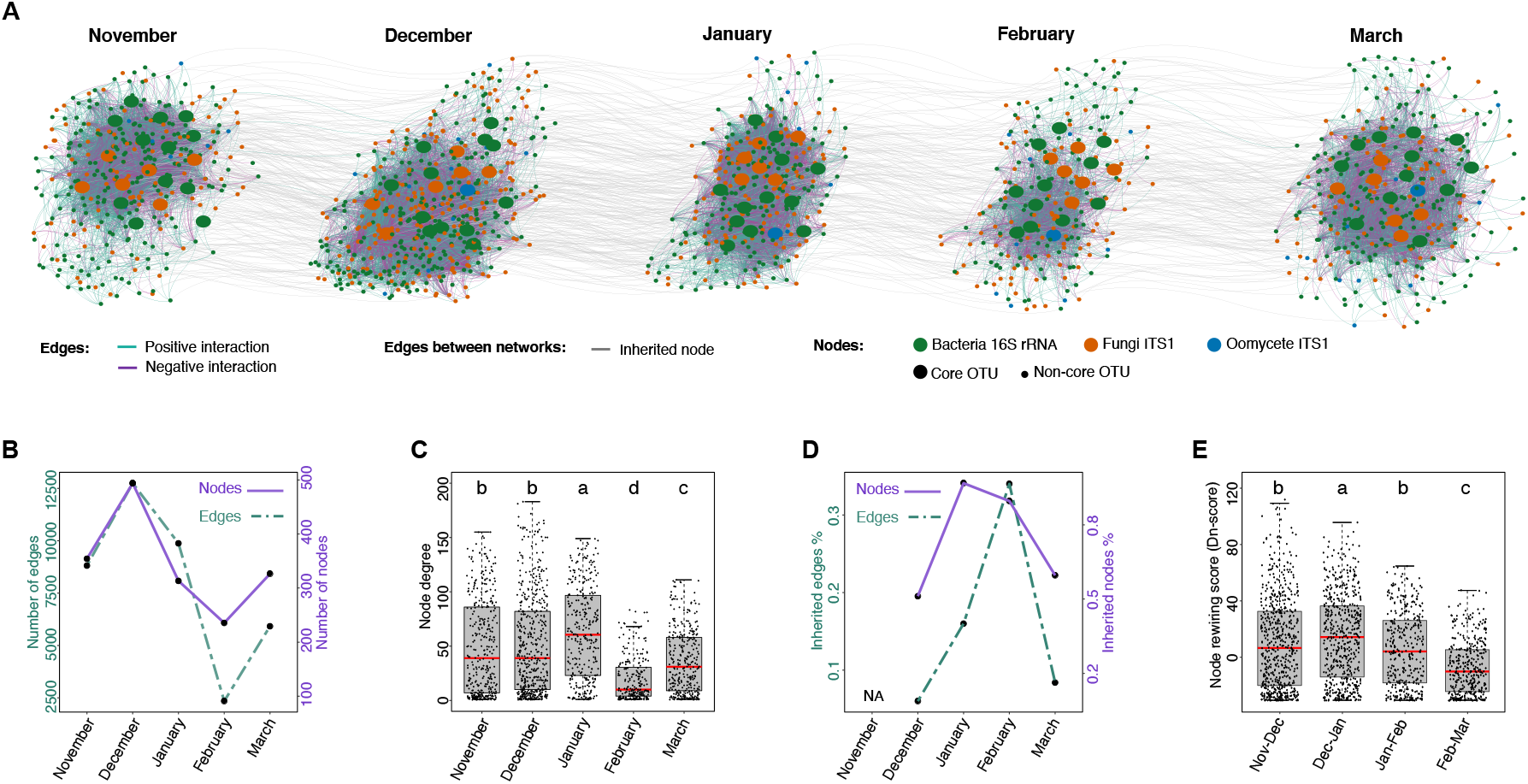
Changes in phyllosphere microbial interaction networks throughout *A. thaliana’s* growing season. **(A)** Data from the three experiments was aggregated to reconstruct coabundance networks for each time point (month) using the SparCC algorithm. Nodes (dots) represent OTUs, edges (colored lines) depict potential positive and negative interactions between OTUs (connections). Nodes from core microbes are indicated. Grey lines connecting networks show nodes conserved from one month network to the next (inherited nodes). **(B)** Number of nodes and edges in each month network **(C)** Percentage of nodes and edges in a given month network which are inherited from (shared with) the previous month network. **(D)** Percentage of edges inherited for a given inherited node. **(E)** Node degree i.e. number of edges per node in each month network. **(F)** Node-rewiring score (Dn-score) calculated in DyNet. For each node, its connected neighbors are compared between two networks (consecutive months) and the changes (rewiring) are quantified. Points represent rewiring scores from single nodes, high values indicate important changes in the node’s connections between the compared networks. Different letters indicate significant differences between conditions (Dunn test, *P* < 0.05).

With the hypothesis that these changes were associated to an increased stability of the network’s structure, we compared networks from consecutive months, recording similarities (inherited nodes/edges) and differences (node rewiring events) between them. Inherited nodes/edges were defined as those shared between consecutive months. The percentage of inherited nodes per network increased from 51% in December to 89% in January and 82% February, meaning the large majority (82 %) of the nodes in the February network were already present in the January network (Fig. 4D). A similar trend was observed for the number of inherited edges, doubling from December (6%) to January (16%) and February (34%). To quantify changes between networks, taking into account the nodes and their connections, we calculated a node-rewiring score for each node in the network. This score reflects the changes in a node’s connections between the compared networks (Dn-score in DyNet) [27]. This analysis revealed that differences between networks tended to decrease through time, with minimum rewiring events between the months of February and March (Dunn test, *P* < 0.05) (Fig. 4E). These results suggest that throughout the beginning of the season (November to February), leaf microbial networks go through a stabilization phase, during which month-to-month changes tend to diminish (increasing numbers of shared nodes and edges, and decreasing node rewiring) as networks exhibit lowering complexity (lower numbers of nodes, edges and connections), reaching minimum levels in February.

### Identifying hubs among core microbes in Arabidopsis leaf microbiota

Time-based microbial networks were analyzed to determine whether potential ‘keystone’ microbes (i.e. hubs - taxa with high betweenness and high closeness centrality) in the leaf microbiota were also highly persistent core microbes. Connectivity analysis on individual month networks revealed few taxa exhibiting hub characteristics (4-10 OTUs, 1-3% of network OTU nodes) and a high turn-over between months, with no taxon systematically identified as hub in every month network (Fig. 5A; Supplementary Table 1). Among the 19 ‘core’ taxa identified previously (Fig. 2) only three bacterial OTUs i.e. Bacillus OTU00012, Oxalobacteraceae OTU00004 and OTU000013 Marmoricola could be identified as hubs, exhibiting high network connectivity in the months of December and February (Fig. 5B; Supplementary Table 1). Blast alignments of 16S rRNA sequences from Oxalobacteraceae OTU00004 further classified this taxon as *Massillia* sp.

**Figure 5.**
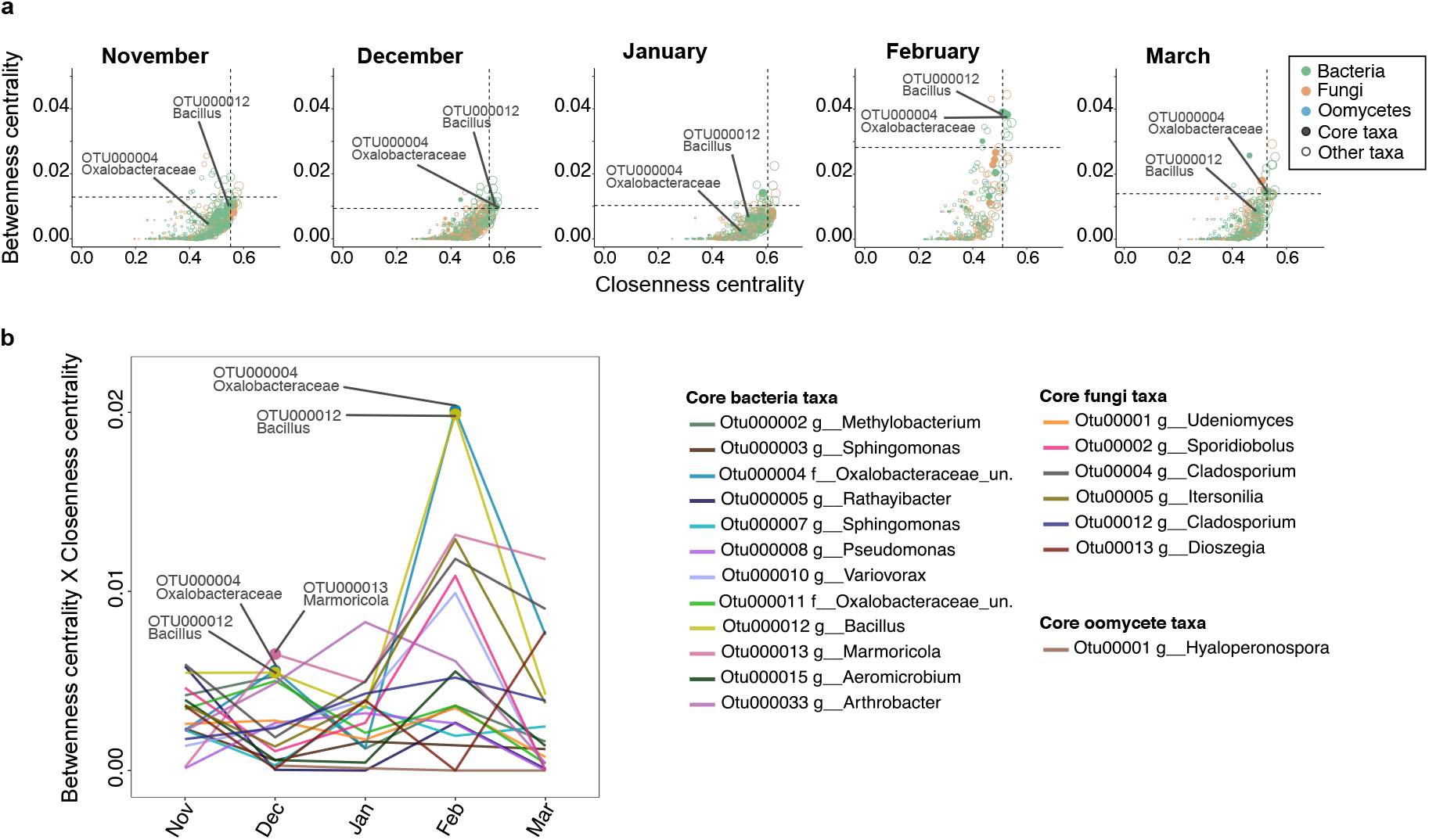
Identification of microbial hubs within *A. thaliana*’s core leaf microbiota. The correlation networks calculated with SparCC (Fig. 3), were used to identify microbial hubs as nodes with high betweenness centrality (i.e. the fraction of shortest paths passing through the given node) and high closeness-centrality (i.e. the average shortest distance from the given node to other nodes). **(A)** Values for single taxa, dotted lines indicate the top 5% values. Circles are colored based on microbial phyla. Circle sizes depict de node’s degree. Closed circles indicate taxa identified as part of the core leaf microbiota. Two core OTUs (12 and 4) are annotated. **(B)** Changes in the connectivity of core taxa. The product of **“**Betweenness centrality × Closeness-centrality” was used to depict monthly changes in the connectivity of core OTUs. Hub taxa are indicated.

As hub identification is highly dependent on network calculation approaches, we repeated these analyses on Spearman-based correlation networks calculated in Co-Net (Fig. S4) with partially similar results. Approximately a third of the OTUs identified as hubs in the SparCC networks were also identified as hubs in the Spearman-based networks (Supplementary Table 1). Notably, this also included Bacillus OTU00012 and Oxalobacteraceae OTU00004, classified as *Massillia* sp. Taken together these results indicate that, with the exception of one *Massillia* and one *Bacillus* lineage, ‘core’ taxa in the Arabidopsis leaf microbiota are not major network hubs and that network hub microbes change over time.

## Discussion

The phyllosphere is a complex microbial habitat due to its direct exposure to a range of abiotic factors --light, humidity and temperature-- that can alter the leaf environment within minutes, hours or days. Furthermore, leaf microbial communities are directly exposed to the arrival of new microbes disseminated by soil particles, water and wind [28]. In this context, key ecological questions are still unanswered: what is the relative importance of environmental filtering versus biotic interactions in shaping community structures and what is the impact of stochasticity [29]? Our limited understanding of the processes behind the assembly and persistence of microbes on leaves under field conditions and their colonization of leaf surfaces constitutes a major drawback for the agricultural usage of plant-beneficial microbes [30]. To address these fundamental questions, we have conducted a long-term experiment to follow month-to-month changes in the composition of Arabidopsis’ leaf microbiota during the early growing season (November to March).

As expected for dynamic ecological systems [31], bacteria, fungi and oomycete leaf-associated communities were highly stochastic, with factors like the sampling time and the plant ecotype explaining only half the variability observed (Fig. S1). Despite high between-experiment variability, robust differences between months were observable for some microbial groups known to be relevant for plant-growth like Peronosporales oomycetes (Fig. S2). *Hyaloperonospora*, the causal agent of downy mildew was by far the most abundant Peronosporales in Arabidopsis leaves, as it has been described for various geographic locations elsewhere [18]. Although our sampled plants exhibited no downy mildew disease symptoms at any time throughout the field experiments, the relative abundance of Peronosporales increased throughout the growing season reaching maximum values in March. This is in agreement with disease dynamics of downy mildew in Brassicaceae known to be favored by cold wet weather, and could indicate that the pathogenic pressure on the plant increases over the early growing season.

The analysis of community dynamics indicates that from November to February leaf microbial communities go through a stabilization phase becoming less diverse, less variable and as microbial networks become less complex (Fig. 3; Fig. 4). This is likely a result of the fact that core microbes become increasingly dominant throughout the season (Fig. 3B). Seasonal dynamics have been described in microbiota associated with plants [11, 12, 32] and animals [33–36], and are thought to be driven partly by environmental cues and perturbations. The fact that in our study microbiota dynamics mirror temperature and rainfall decreases associated with winter (Fig. 1), lead us to hypothesize that climatic conditions might be driving the observed leaf microbiota dynamics, maybe via the selection of cold-resistant microorganisms. Indeed a strong “winter effect” on microbial communities has been observed in a diversity of environments including the bee’s gut [36], lake water [37] and air [38]. We hypothesize that winter conditions might apply a strong selective filter causing leaf microbial communities to reduce in complexity. Longer experiments are needed to determine if different dynamics would be observed in later stages e.g. during spring.

Microbes with a stable presence in *Arabidopsis* leaves (core taxa; Fig. 2) accounted for only 0.62 % of all detected leaf-taxa, indicating a high turnover in leaf microbial communities. Core taxa include putative plant pathogens like Hyaloperonospora and Cladosporium [39, 40] but at the same time taxa encompassing plant beneficial microorganisms like Sphingomonas and Variovorax, which could explain the asymptomatic state of the sampled plants. Leaf-inhabiting Sphingomonas bacteria have been shown to protect Arabidopsis from bacterial pathogens [2] and are hypothesized to participate in plant disease resistance against root fungal pathogens. Variovorax strains have been shown to modulate plant hormonal balance by degrading auxins thus promoting plant growth under stress conditions [41]. But not only bacteria have been reported to interfere with plant hormone levels, there have been reports of yeasts on *A. thaliana* capable of producing auxin-like indolic compounds [42]. We have identified four basidiomycete yeast taxa (*Udeniomyces, Sporidiobolus, Itersonilia* and *Dioszegia*) as systematic colonizers of Arabidopsis leaves. Although little is known about the associations between these yeasts and Arabidopsis, a recent study on a leaf basidiomycete yeast (*Moesziomyces bullatus*) suggests they can play important roles in plant protection from pathogenic oomycetes through protein effectors [43]. While previous studies have reported on the prevalence of some of the identified core taxa on Arabidopsis’ leaves [16, 44, 45], here we show these associations are time-stable persisting throughout the plant’s life and between plant generations, suggesting some level of adaptation to the leaf niche on the microbial side or even possible co-evolution between these microbes, as well as with the host plant.

Microbe-microbe interactions participate in the structuring of microbial communities with certain microbes --hub and keystone microbes-- playing central roles [46]. We hypothesized that high connectivity within leaf microbial networks might explain the persistence of the identified core taxa. However, in contrast to our hypothesis, the connectivity level (hubness) of individual core taxa was highly variable from month to month, with no taxon maintaining high connectivity levels throughout the entirety of the growing season (Fig. 5). This indicates that high connectivity is not a prerequisite for high prevalence in the leaf microbiota as core taxa are not necessarily network hubs [13]. Nevertheless two microbes among the leaf core taxa, within the *Bacillus* and *Massillia* lineages, deviate from this rule and have been identified as hubs. Interestingly, in the month of February when leaf microbial communities displayed the lowest levels of complexity, both *Bacillus* and *Massillia*, reached maximum connectivity levels within leaf microbial networks (Fig. 5), while their relative abundances on leaves remained stable (Fig. 2). It is tempting to speculate that there might be a functional link between these hubs and community stability. Indeed, it has been shown that highly connected microbes can be good predictors of the stability of microbial communities [47]. In the future, experimental evidence will be needed to improve predictions and to determine if (and how) hub-removal affects the stability of microbial communities, over time.

Taken together our results show that despite a high level of stochasticity leaf microbial communities exhibit detectable time patterns, with stable and unstable components. Although this study is purely descriptive it opens a new field of research on time-informed community dynamics in natural host-associated microbiomes. In the long term, these types of studies could make it possible to model, predict and drive microbial communities to desired states.

## Author Contributions

The project was initiated by S.K., M.A. and E.K. S.K., M.A. and A.M. conducted the experiments with help from A.P. J.A. and M.M. analyzed the data and J.A., M.M. and E.K. wrote the manuscript with contributions from all authors.

## Acknowledgements

This work was supported by the Max-Planck Gesellschaft, the University of Tübingen, the European Research Council (ERC) under the DeCoCt research program (grant agreement: ERC-2018-COG 820124), the Cluster of Excellence on Plant Sciences (CEPLAS; Exc 1028) and the SPP 2125 DECRyPT program from the DFG.

## Competing Interests

Authors declare no competing financial interests in relation to the work described.

## Supplementary figures

**Supplementary figure 1.**
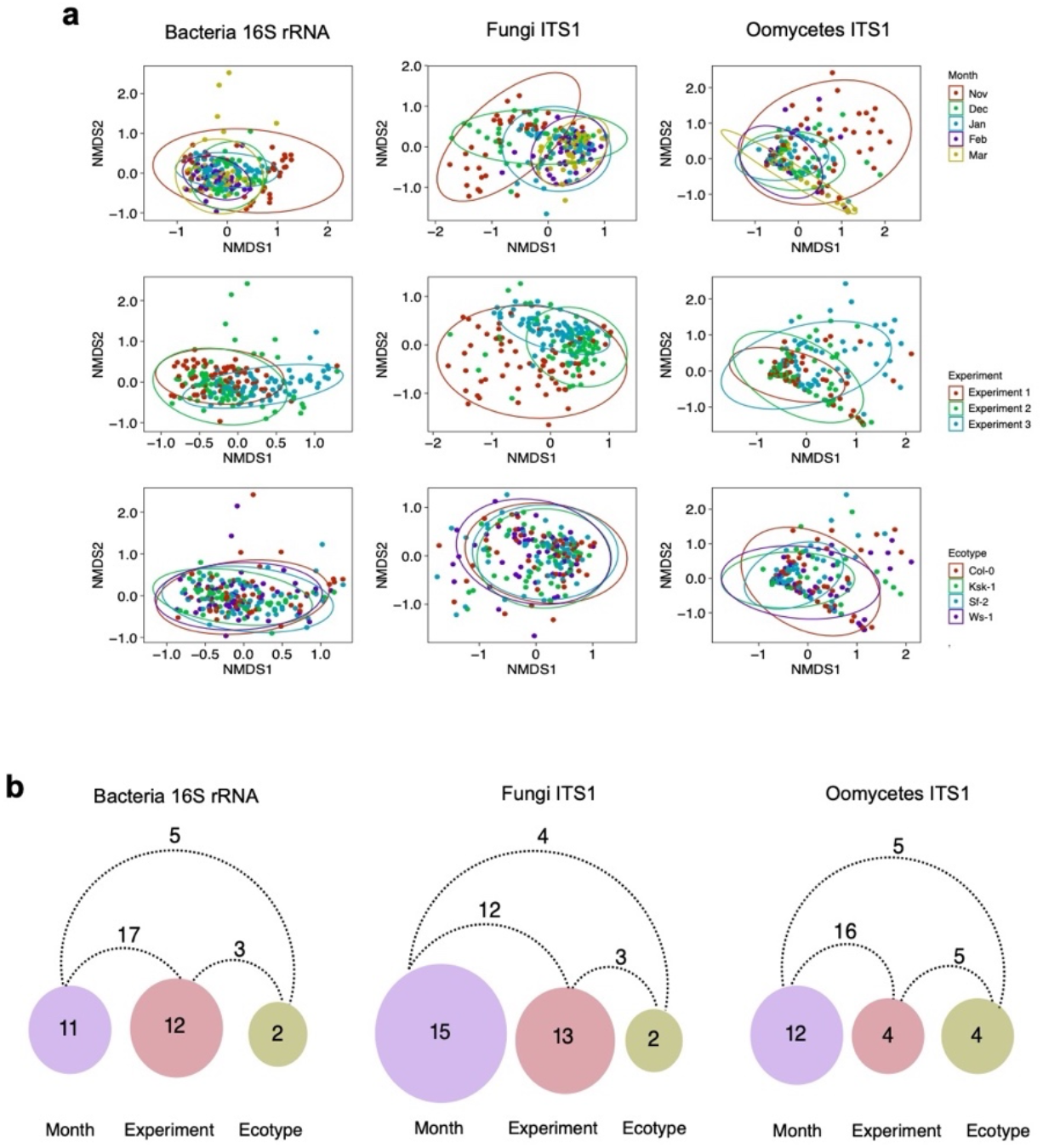
Multivariate analysis on factors structuring leaf microbial communities. **(A)** Non-metric multidimensional scaling ordination (NMDS) on Bray-Curtis dissimilarities between samples grouped by ‘month’, ‘experiment’ or ‘ecotype’. **(B)** Circles depict the percentage of variance explained by factors ‘month’, ‘experiment’ and ‘ecotype’, connecting lines depict the percentage of variance explained by interactions between factors. A PerMANOVA analysis on Bray-Curtis distances was conducted using the Adonis function in Vegan. Only significant effects are shown (permutations 10000, *P* < 0.05, explanatory categorical variables: Experiment x Month x Ecotype).

**Supplementary figure 2.**
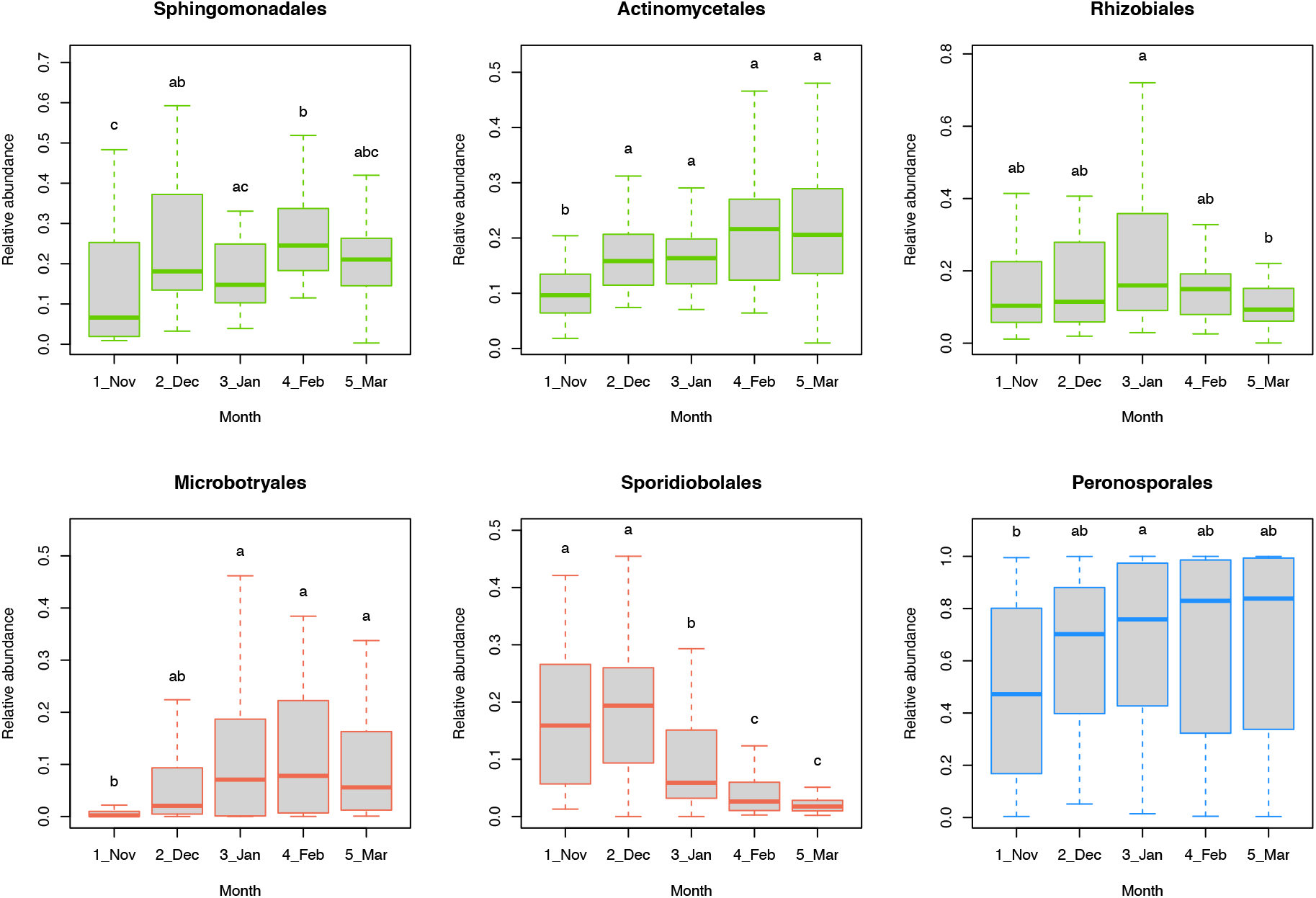
Temporal changes in high abundance microbial taxa colonizing *A. thaliana*’s leaves. Boxplots show the relative abundance of the bacterial (green), fungal (orange) and oomycete orders in single samples aggregated by ‘month’. Whiskers depict the dispersion of the data (1.5 x interquartile range) and different letters indicate significant differences between months (ANOVA followed by Tukey’s HSD, *P* < 0.05).

**Supplementary figure 3.**
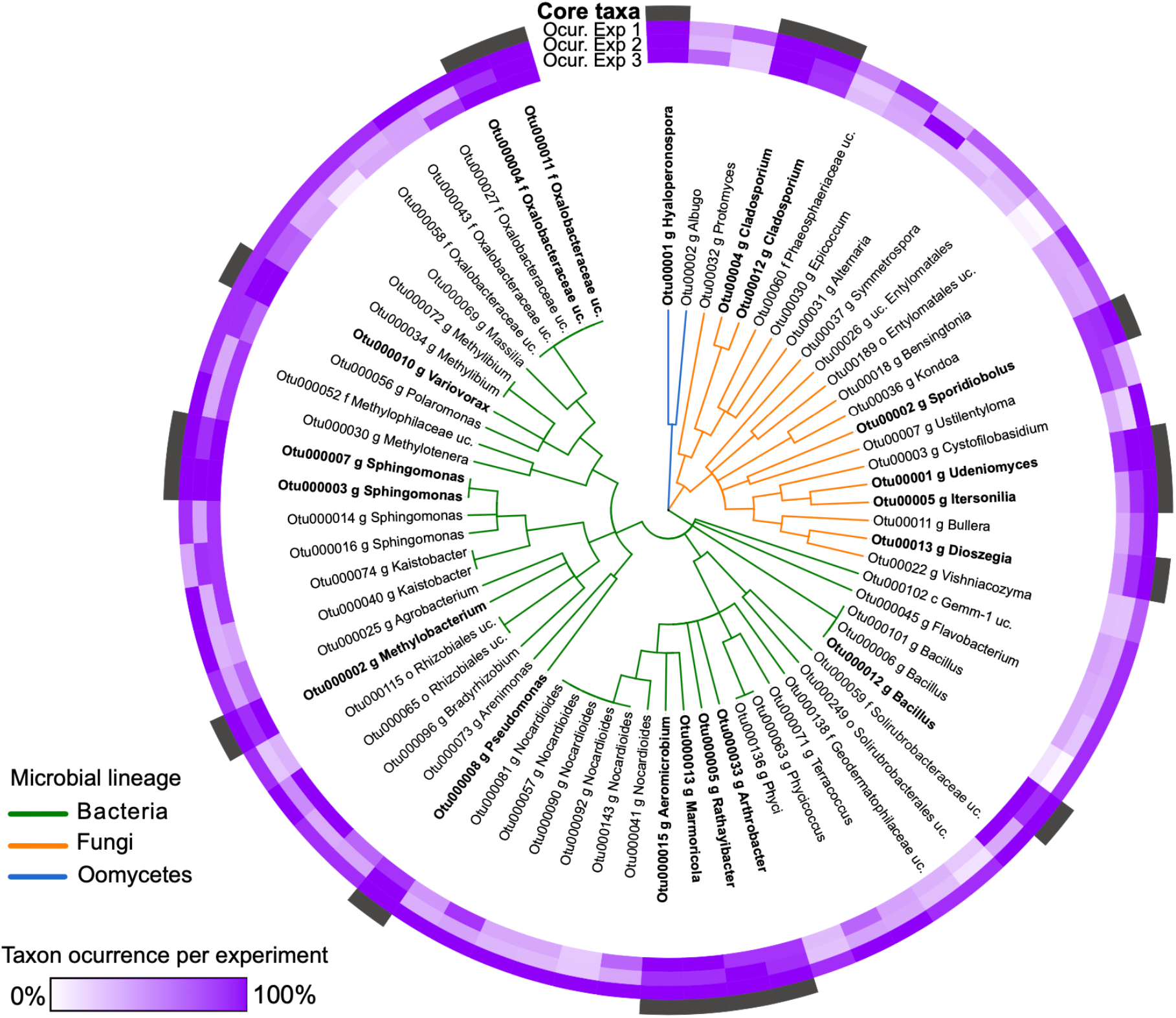
Identification of persistent core microbial taxa. Persistent core taxa were identified as OTUs showing high-occurrence (≥ 95% for fungi and oomycete, ≥ 98% for bacteria) in each of the three experiments analyzed. Purple inner rings depict OTU occurrence within each year while the outer black ring denotes OTUs identified as “core” (names highlighted in bold) (See Fig. 3).

**Supplementary figure 4.**
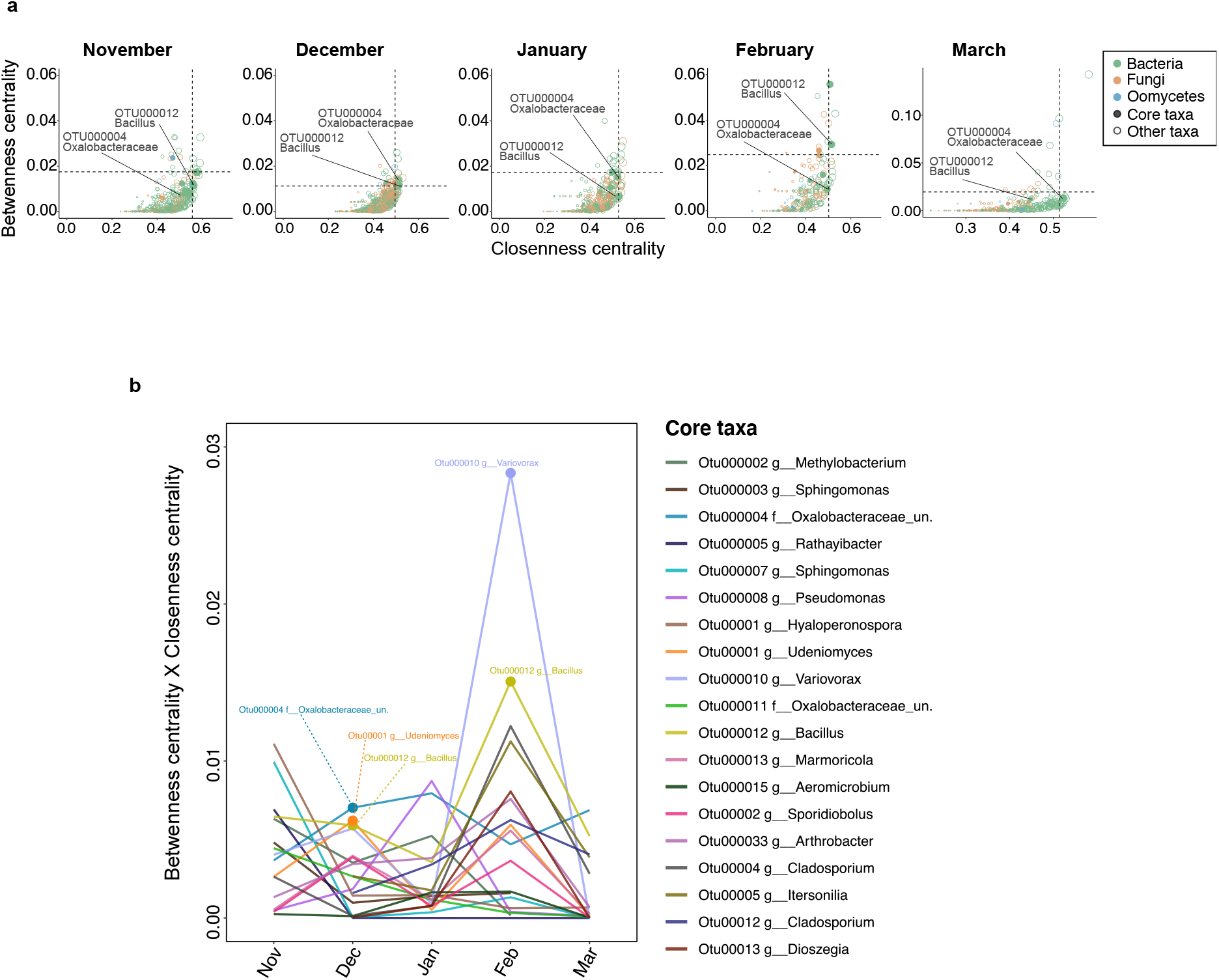
Identification of microbial hubs within the leaf microbiota of *A. thaliana* (Co-Net-based networks). **(A)** Co-Net-based Month networks were used to identify microbial hubs as nodes with high betweenness centrality (i.e. the fraction of shortest paths passing through the given node) and high closeness-centrality (i.e. the average shortest distance from the given node to other nodes). In each graph, dotted lines indicate the top 5% values. Circles are colored based on microbial phyla. Circle sizes depict de node’s degree. Closed circles indicate taxa identified as part of the core leaf microbiota. Two core otus (12 and 4) are highlighted. **(B)** Changes in the connectivity of core taxa. The product of “Betweenness centrality × Closeness-centrality” was used to depict monthly changes in the connectivity of core OTUs. Identified hub taxa (panel A) are indicated.

## Notes

### Competing Interest Statement

The authors have declared no competing interest.

### Summary of Updates

Corrections in author name and affiliations

